# Naa80 is required for actin N-terminal acetylation and normal hearing in zebrafish

**DOI:** 10.1101/2024.03.17.585386

**Authors:** Rasmus Ree, Sheng-Jia Lin, Lars Ole Sti Dahl, Kevin Huang, Cassidy Petree, Gaurav K. Varshney, Thomas Arnesen

**Affiliations:** Department of Biomedicine, University of Bergen, Norway; NORCE Climate & Environment - NORCE Norwegian Research Centre, Bergen, Norway; Genes & Human Disease Research Program, Oklahoma Medical Research Foundation, Oklahoma City, OK, USA; Virology Unit, Faculty of Veterinary Medicine, Norwegian University of Life Sciences; Department of Surgery, Haukeland University Hospital, Norway

**Keywords:** Actin, N-terminal acetylation, zebrafish, NAA80, cytoskeleton, hearing loss

## Abstract

Actin is a key component of the cytoskeleton of eukaryotic cells and is involved in numerous cellular functions. In animal cells, actins are uniquely N-terminally processed by a dedicated enzyme machinery to generate their mature acidic and acetylated forms. The final step of this maturation process involves N-terminal acetylation, a reaction catalyzed by NAA80 in humans. In human cell lines, N-terminal acetylation of actin plays a crucial role in maintaining normal cytoskeletal dynamics and cell motility. The physiological impact of actin N-terminal acetylation remains to be defined. Here, we developed a zebrafish *naa80* knockout model and established that zNaa80 acetylates both muscle and non-muscle actins *in vivo*. Our *in vitro* investigation of purified zNaa80 unveiled a clear preference for acetylating N-termini derived from actins. Interestingly, zebrafish lacking actin N-terminal acetylation were viable and exhibited normal development, morphology and behaviour. In contrast, human individuals carrying pathogenic actin variants may present with hypotonia and hearing impairment. While zebrafish depleted for *naa80* did not display any obvious muscle defects or abnormal muscle tissue, we found that they have abnormal inner ear development such as small otoliths and impaired response to sound stimuli. In sum, we have defined that zebrafish Naa80 N-terminally acetylates actins *in vitro* and *in vivo* and that actin N-terminal acetylation is essential for normal hearing *in vivo*.

## Introduction

Actin is the most abundant protein in animal cells and participates in a great variety of cellular functions. Actin filaments maintain the shape and rigidity of a cell and provides the scaffold for myosin-driven movements which power protrusion formation and muscle contraction (*1, 2*). Actin-myosin powered cell movements are prevalent and of great importance during development (see for example (*3*)). Actin function is regulated through several different mechanisms, among these post-translational modifications (PTMs) and actin-binding proteins (ABPs) (*4*). In the animal kingdom, actin undergoes a unique N-terminal (Nt) maturation process involving a final Nt-acetylation step(*5*–*7*). The enzyme responsible for this final maturation step, NAA80, was only recently identified (*8*–*10*). Nt-acetylation of the cytoplasmic actin isoforms (β-actin and γ-actin in humans) is very prevalent, with around 100% of the actin so modified in HAP1 and other mammalian cells (*8*). This acetylation is lost in *NAA80*-KO cells (*8, 11*). Non-acetylated actin polymerizes and participates in functional F-actin networks. However, it is slower to polymerize and to depolymerize *in vitro*, in sum yielding an increased filamentous to globular (G/F) actin ratio compared to wild-type cells (*8*). *NAA80*-KO cells show increased rates of migration, increased cell size, Golgi fragmentation and increased numbers of filopodia and lamellipodia (*8, 12, 13*). NAA80 has a conserved polyproline loop which is required for its association with the actin-binding protein PFN2 (*14, 15*). The NAA80-PFN2 association facilitates efficient post-translational Nt-acetylation of G-actin monomers before these are incorporated into F-actin.

The first description of patients with a pathogenic *NAA80* variant was published recently (*16*). Two siblings presented with progressive high-frequency hearing loss, craniofacial dysmorphisms, developmental delay and mild muscle weakness of both the trunk and the limbs. Sequencing showed that the patients were homozygous for a c.389T>C (Leu130Pro) variant in *NAA80* which resulted in destabilization of the NAA80 protein and a reduction of actin Nt-acetylation to approximately 50% of the level in control cells. Several of the observed symptoms overlapped with known symptoms of actin loss-of-function variants.

The NAA80 knockout phenotype in single cells is well characterized, but there currently are no loss-of-function animal models of *NAA80*. To remedy this, we have studied the function of Naa80 in *Danio rerio* (zebrafish) by investigating its knockout phenotype and characterizing the substrate specificity of Naa80. We find that zebrafish Naa80 Nt-acetylates actin-type N-termini and that *naa80* -/- zebrafish lack actin Nt-acetylation of cytoplasmic and muscle actins. The *naa80* -/- zebrafish are apparently morphologically normal and thrive normally.

However, one of the knockout lines did not yield any eggs, raising the possibility that Nt-acetylation of actin has a function in egg-laying. Furthermore, *naa80* KO larvae have smaller otoliths, fewer hair cell bundles and reduced viability of hair cells in the inner ear. This was accompanied by a decreased startle response, consistent with a hearing abnormality.

## Results

### A *Danio rerio* Naa80 orthologue displays Nt-acetyltransferase activity towards acidic, actin-type N-terminal peptides *in vitro*

Like other animals, zebrafish have several actins which differ in their Nt-sequences. The actin Nt-sequences have subtle differences but are invariably acidic (Fig. 1). Class I actins (cytoplasmic 1 and cytoplasmic 2) have a stretch of 3 acidic amino acids at the N-terminus following the initiator methionine (iMet), while class II actins (alpha-cardiac 1a and 1b, alpha-skeletal 1a and 1b, and alpha-smooth) have iMet and cysteine residues N-terminal to 3-4 acidic residues.

**Figure 1:**
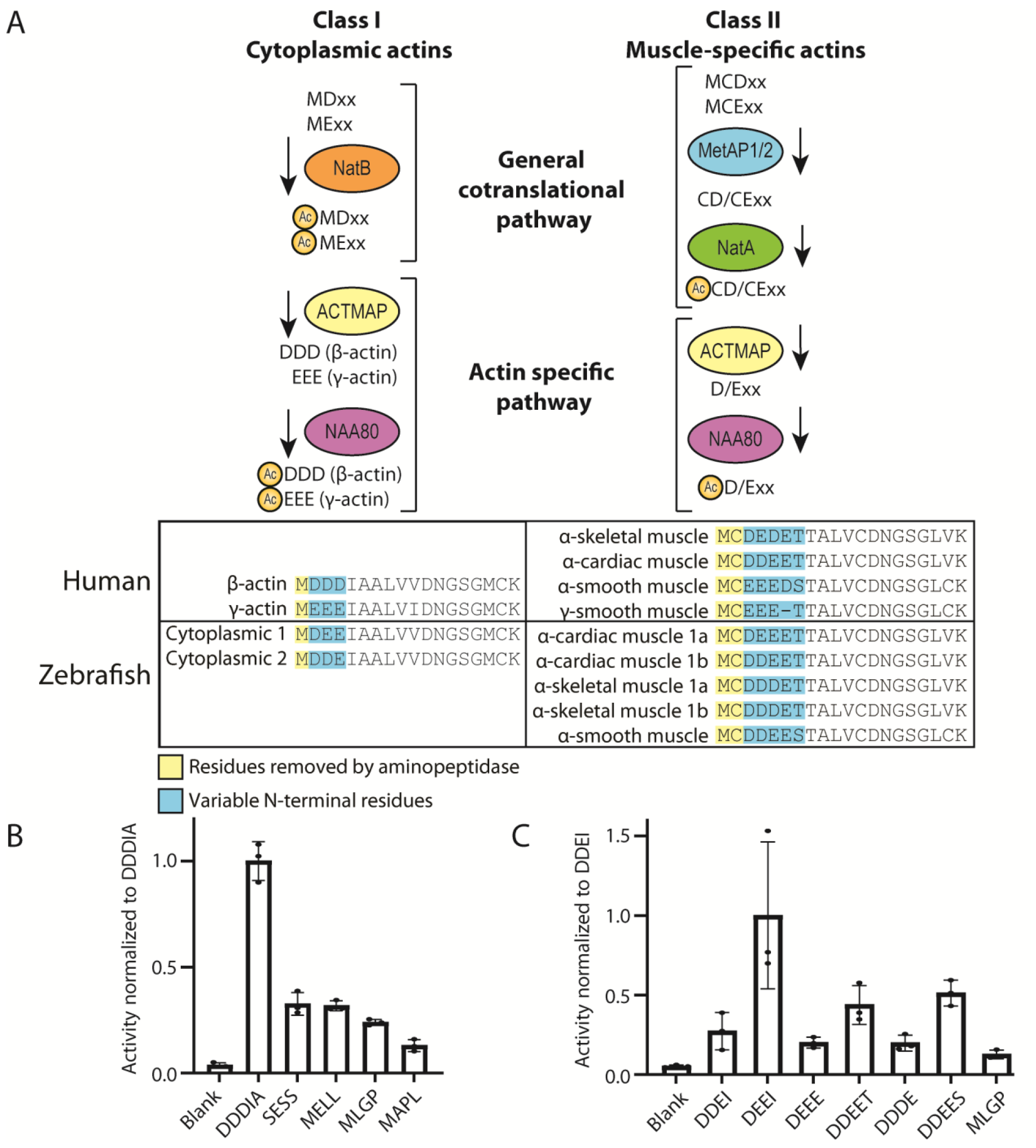
Actin N-terminal maturation scheme and the *in vitro* acetyltransferase activity of *Danio rerio* E7FBQ5/Naa80 towards actin-type N-termini. A) Overview of actin N-terminal processing leading up to Nt-acetylation. Class I actins are first co-translationally Nt-acetylated by NatB, then actin methionine aminopeptidase (ACTMAP) removes the acetylated Met, before NAA80 acts on the acidic neo-N-terminus. Class II actins are processed by MetAP1/2 before Nt-acetylation by NatA and removal of an acetylated Cys by ACTMAP before Nt-acetylation by NAA80. In both humans and zebrafish, actin is conserved with most of the amino acid differences located to the N-terminus. B) Acetylation assays with purified MBP-E7FBQ5/zNaa80 and different peptides derived from human proteins. C) Acetylation assays with purified MBP-E7FBQ5/zNaa80 and different peptides derived from zebrafish actins.

The class I actins are presumed to be co-translationally acetylated by NatB (*17*) before the Nt-Ac-Met is removed by ACTMAP, the recently discovered actin maturation protease (*18*). This exposes the neo-N-terminus at Asp2/Glu2 and allows post-translational Nt-acetylation by NAA80 (*8*). The class II actins (Met-Cys-Asp/Glu-…) are presumed to be substrates of MetAP1/2, removing the iMet and exposing the neo-N-terminus at Cys2 (*19*). This is then potentially Nt-acetylated by NatA (co-translationally) before the Cys2 is removed by ACTMAP (*18*), exposing the Nt-amine group of acidic Asp3/Glu3 as the acceptor of NAA80-mediated acetylation (*20*).

To identify the zebrafish ortholog of NAA80 we performed a protein BLAST (*21*) search using the human NAA80 protein sequence as a query. The top-ranked hit is annotated in Uniprot (E7FBQ5) as N-alpha-acetyltransferase 80, NatH catalytic subunit, because of its high sequence similarity with NAA80 from other organisms (*10*). We cloned the cDNA encoding this protein (Refseq: XM_005167127.3) and expressed it in *Escherichia coli* BL21* cells with maltose-binding protein (MBP) and His-tags to enable purification. After a two-stage purification process we obtained apparently homogenous MBP-E7FBQ5/zNaa80.

Purified zNaa80 was used in an acetylation assay to determine its activity and potential substrate preference. The enzyme was tested against peptides with different N-terminal sequences, and the activity readout is disintegrations per minute (DPM) (*22*) normalized to the peptide with the highest activity (Fig. 1B-C). We initially tested several peptides representing different NAT substrate classes (Fig. 1B). NAA80 is defined by the activity towards acidic N-termini without the initiator methionine, represented by the N-terminus DDDIA, which is derived from human β-actin, and by EEED, DDEE, and DEDE, which are derived from human alpha-smooth muscle, alpha-cardiac muscle, and alpha-skeletal muscle respectively. SESS is derived from a NatA substrate (*23*), MELL derived from a classical NatB substrate (*17*), MLGP is derived from a NatC/NatE substrate (*24*), and MAPL represents a NatF substrate (*25*). The activity profiles of E7FBQ5, preferring acidic actin N-terminal peptides, strongly suggest that this protein indeed represents zNaa80.

### Dynamic expression of *naa80* mRNA at different developmental stages and adult tissues

To gain insight into the possible function of *naa80*, we employed whole-mount *in situ* hybridization (WISH) (Figure 2A-D) and RT-qPCR (Figure 2E) at different developmental stages to unveil the spatial and temporal *naa80* mRNA expression pattern. Remarkably, we detected *naa80* expression as early as 1-hour post-fertilization (hpf) (Figure 2A), suggesting *naa80* mRNA maternal distribution. By 24 hpf, *naa80* mRNA exhibited ubiquitous expression, encompassing the central nervous system, eyes, otic vesicles, and trunk muscles (Figure 2B and B’). Subsequently, at 48 and 72 hpf, *naa80* expression became restricted to the brain, eyes, and pectoral fin bud (Figure 2C, C’, and D). The RT-qPCR results supported the presence of *naa80* expression throughout embryonic development (Figure 2E).

**Figure 2:**
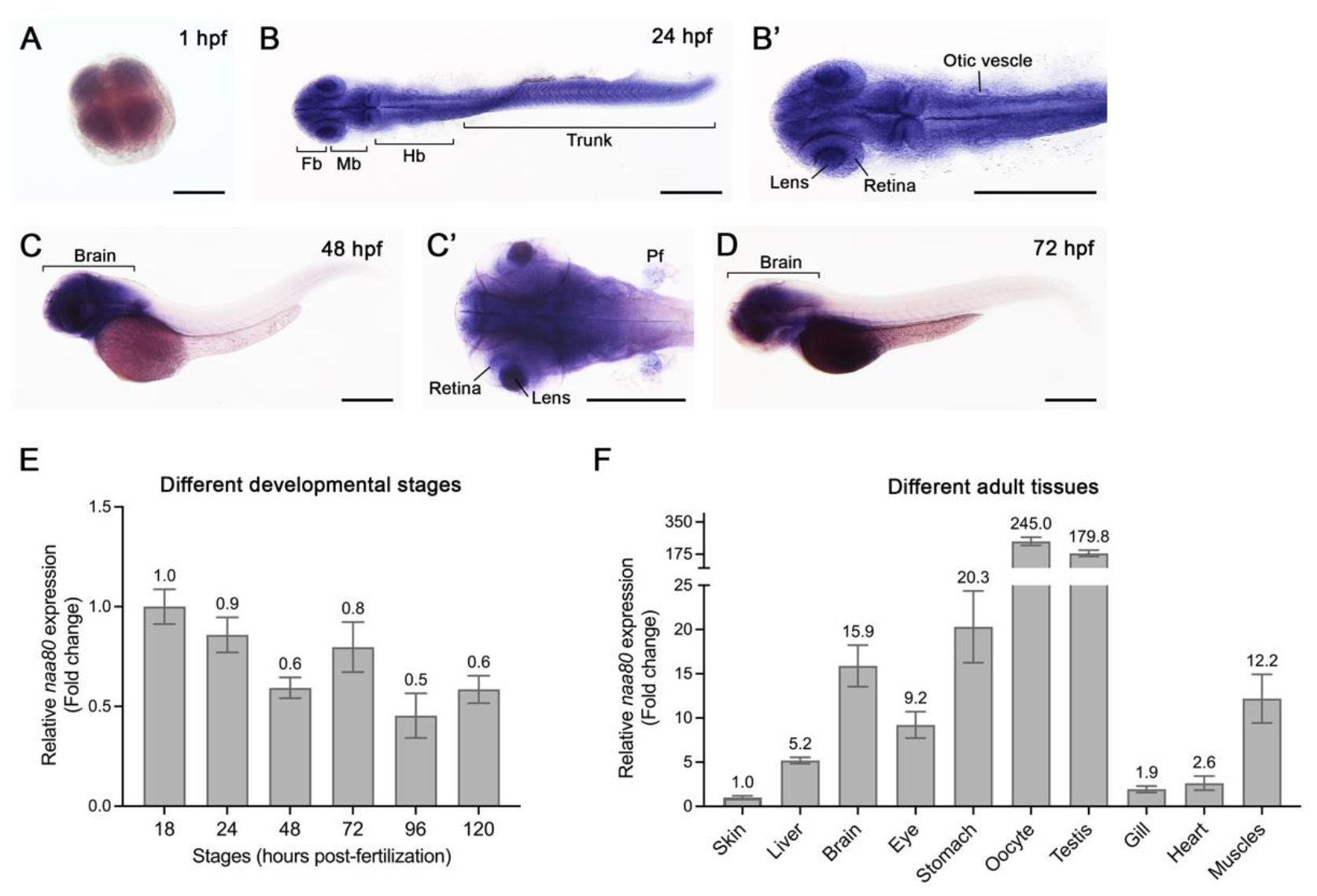
The spatiotemporal expression of *naa80* mRNA by WISH and RT-qPCR. WISH was performed at different developmental stages of zebrafish embryos at 1 **(A)**, 24 **(B, B’)**, 48 **(C, C’)** and 72 **(D)** hpf. RT-qPCR at different embryonic stages **(E)** and adult tissues **(F)**. For (A, B, B’ and C’), dorsal view and anterior to the left. For (C, D), lateral view and anterior to the left. Fb, forebrain. Mb, midbrain. Hb, hindbrain. Pf, pectoral fin bud. Scale bar = 0.3 mm. For (E, F), each timepoint included biological triplicates as well as technical triplicates. The expression levels were first normalized to the *18S* housekeeping gene and the expression levels were compared to 18 hpf embryos (E) or compared to liver (F). Data shown are mean ± SD and mean values were shown on each bar.

Furthermore, we conducted RT-qPCR analysis on various tissues from one-year-old adult animal. The results unveiled *naa80* expression in most tissues, with notable enrichment in the kidney, stomach, oocytes, testis, brain, and skeletal muscles (Figure 2F). This enrichment suggests a potential role for *naa80* in these tissues. In summary, *naa80* mRNA exhibits ubiquitous expression during embryonic development, gradually becoming more confined to the brain, and it displays relatively higher levels in kidney and reproductive tissues during adulthood.

### *naa80* -/- zebrafish are free from gross morphological phenotypes and develop normally

To study the *in vivo* function of zNaa80, we had two zebrafish knockout lines for the putative *naa80* gene generated. The knockout alleles were verified by PCR using *naa80*-specific primers (see Table 1) followed by either Big Dye sequencing or agarose gel electrophoresis. Knockout fish were morphologically normal and developed normally (Figure 3). Male *naa80* 13del+/- fish were significantly shorter than controls (p < 0.05); however, *naa80* 13del-/- fish were not significantly shorter so the *naa80* genotype is not likely to be the cause of this. In all other comparisons, body weight and length were not different from clutch mates, and we did not find any morphological abnormalities specific to *naa80* +/- or -/- fish. We attempted to obtain *naa80*-/- eggs by crossing *naa80* 5del/1in -/- and *naa80* 13del -/- fish in the F1 generation. However, this did not yield any eggs. For this reason, the F2 generation fish were obtained by incrossing *naa80*+/- fish and genotyping the offspring once they reached maturity. In F2, we performed more systematic mating tests to investigate whether this could be a feature of the *naa80*-/- phenotype. Several incrossings of *naa80* 5bp del/1bp ins -/- were performed, but no eggs were obtained. However, *naa80* 13bp del +/- females crossed with *naa80* 13bp del -/- males did yield embryos which were viable and with no apparent phenotype. Likewise, Spotty wild-type females crossed with *naa80* 5del/1in -/- males laid eggs, yielding viable embryos. Crossing *naa80* 5bp del/1bp ins -/- females with Spotty wild-type (SWT) males yielded no eggs.

**Figure 3:**
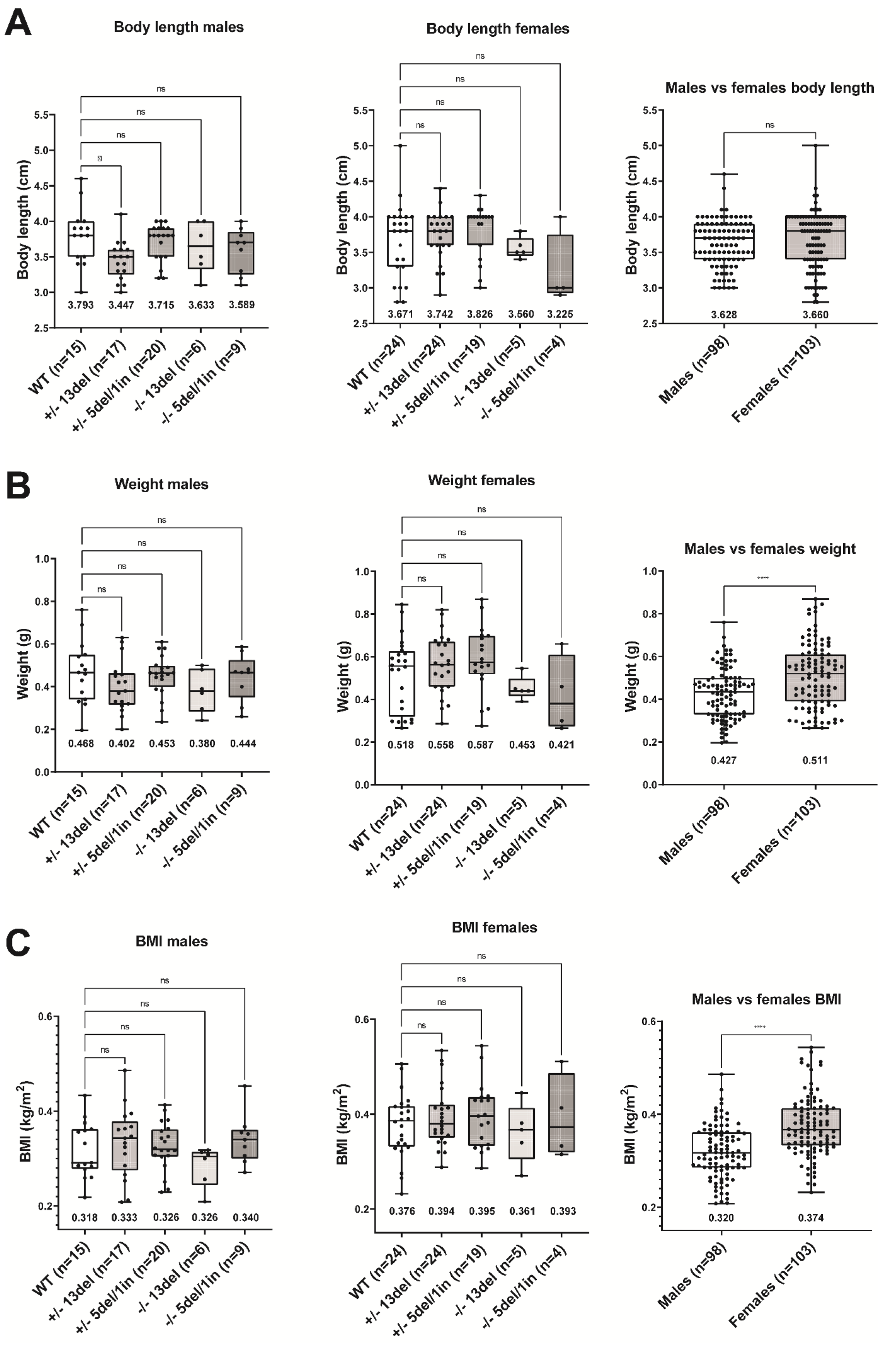
Body length and weight are not affected by *naa80* genotype. Adult zebrafish with the indicated genotype were weighed and measured from rostrum to caudal tip. A one-way ANOVA test followed up with a Dunnett’s test was performed for the genotype comparison segregated by sex, where the +/+ values was selected as the control mean. *: p < 0.05, nonsignificant (ns): p > 0.05. Unpaired t-test was performed for male vs female comparison. ****: p < 0,0001, ns: p > 0.05. The mean value for each dataset is indicated beneath the boxplot. (A) Body length values for males and females. (B) Measured weight of males and females with indicated sample size and mean. (C) The BMI for males and females was calculated by using the formula 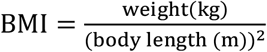.

However, incrossing *naa80* 13bp del -/- fish in F3 resulted in several eggs which developed normally throughout the larval stage. This demonstrates that neither maternal *naa80* nor maternal Nt-acetylated actin are necessary for normal development pre-mid blastula transition. Further, we cannot conclude that *naa80* -/- females are infertile. However, fertility or egg-laying may still be affected, as we were never able to obtain eggs from *naa80* 5bp del/1bp ins -/- females.

### Impaired N-terminal acetylation of cytoplasmic and muscle actins in *naa80* -/- zebrafish

In order to check the Nt-acetylation status of actins in *naa80* KO zebrafish, skeletal muscle and heart tissue were dissected and lysed before soluble proteins were trypsinized and analyzed by LC/MS. We identified substantially higher numbers of peptides and proteins for the heart and skeletal muscle samples, as well as more actin N-terminal peptides and more actin peptides in general. There were no Nt-acetylated actins identified in most *naa80* -/- samples; however, a small amount of Nt-acetylated cytoplasmic actin 2 (Ac-DDEI…) was identified in of the *naa80* 5del/1in -/- heart samples (Figure 4). This could be residual Nt-acetylation by Naa10 or another NAT (*26, 27*), or it could be carryover or contamination from a different sample. Nevertheless, virtually all Nt-acetylation appears to be lost in *naa80* -/- fish.

**Figure 4:**
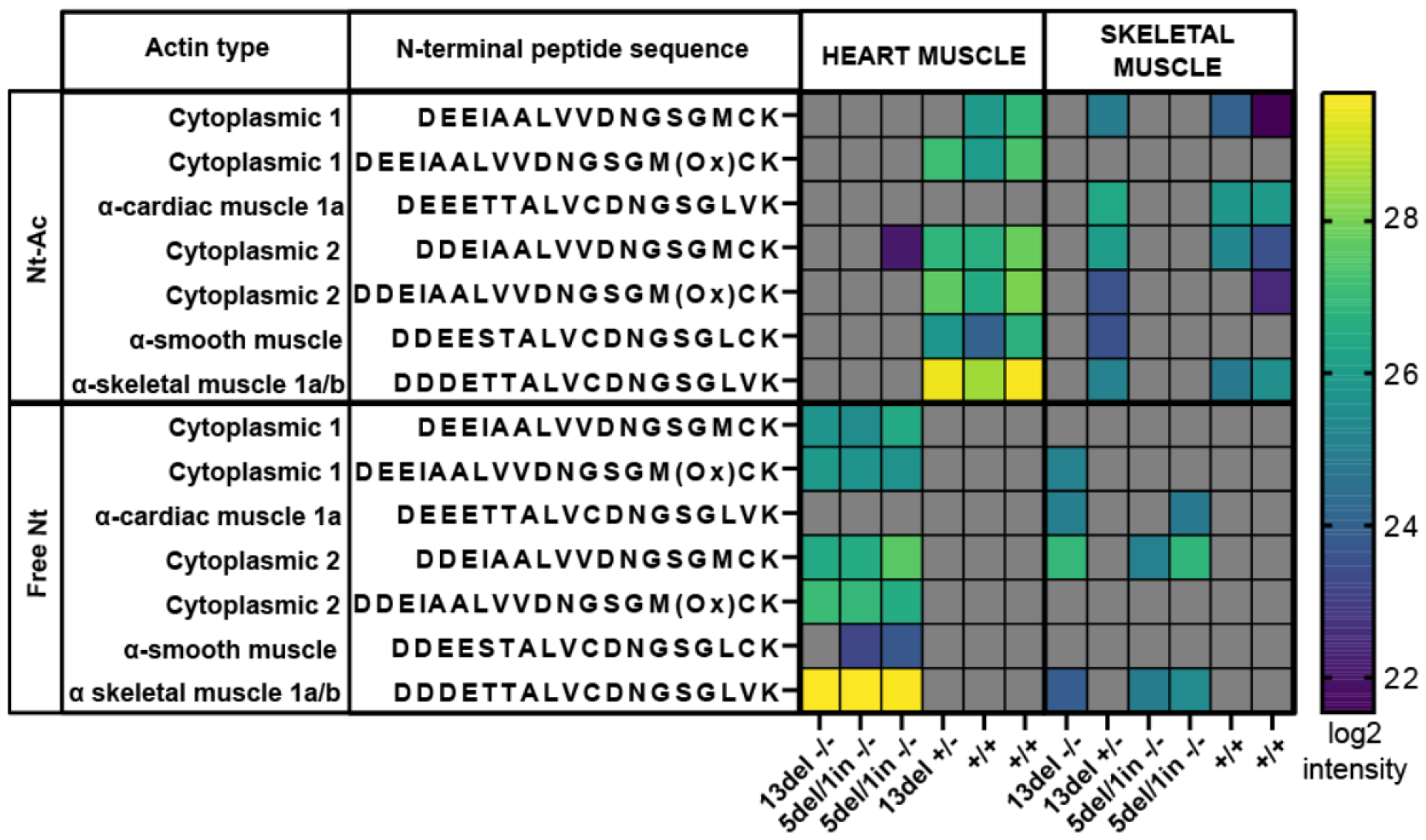
Defective *in vivo* actin Nt-acetylation in *naa80*-/- zebrafish. Mass spectrometry measurements of actin N-terminal peptides in cardiac muscle or skeletal muscle from the indicated genotypes. Log2 intensity of the peptides is reported. Grey: not identified in the sample.

For the *naa80* +/+ and +/-samples we identified Nt-acetylated actin N-termini and very little to no unacetylated actin N-termini. This is in line with earlier studies which show a high degree of actin Nt-acetylation (*8, 11*). We gathered from this that one functional allele of *naa80* is sufficient for complete actin Nt-acetylation. In contrast, we found that in the *naa80* -/- samples from both knockout lines, there is little to no detectable Nt-acetylated actin (Figure 4). We found instead unacetylated actin N-termini, as well as a comparable number of actin non-terminal peptides in *naa80* -/- samples, indicating that this is not due to underrepresentation of actins in the analyzed proteome (Table S2, listing all modified peptides). We thus conclude that Naa80 is the enzyme responsible for class I and class II actin Nt-acetylation in zebrafish, like it is in humans.

### *naa80*-/- F0 knockout (KO) zebrafish displayed hearing-related defects

A previous study indicated that human individuals with biallelic missense variants in *NAA80* exhibit decreased actin Nt-acetylation and increased polymerized actin, potentially contributing to high-frequency hearing loss (*16*). However, no direct hearing-related assessment has been conducted in genetic mutant models. Specifically, how aberrantly acetylated actin leads to the hearing loss phenotype *in vivo* remains unclear. The sensory patches in the inner ear of zebrafish are the maculae and the cristae (*28*). The maculae are located in the otolith organs (utricle and saccule), and the cristae are located in the semicircular canals. Cristae, found in the semicircular canals, detect head movements, and help maintain balance by bending sensory hair cells in response to fluid movement. Maculae, located in the otolith organs, sense linear acceleration and gravity by detecting shifts in otoliths that bend stereocilia of hair cells. The hair cells located in the inner ear’s three cristae and two maculae, as well as in the lateral line neuromast of zebrafish larvae, provide an excellent model for investigating hearing mechanisms. These cells share functional similarities with human inner ear hair cells and are easily accessible for assessment (*29, 30*) (Figure 5A). Actin is essential for the development and function of the inner ear hair cell, including shaping the hair cell, developing stereocilia bundles, and transporting vesicles. To investigate whether the loss of Naa80 contributes to the hearing function and to overcome the fertility difficulties observed in stable mutant lines, we utilized the transient CRISPR method, which efficiently generates null mutants in the founder generation (referred to as knockout or KO hereafter to distinguish them from stable mutants)(*31*). Similar to stable genetic naa80 mutants, we did not detect any evident gross morphological phenotypes in *naa80* KOs. However, we did observe a reduction in otolith size compared to controls following Cas9 protein injection (Figure 5B). Immunohistochemistry analysis utilizing acetylated tubulin (Ac-tub, for kinocilia staining) and phalloidin (staining F-actin, for stereocilia staining) (Figure 5A) indicated that KOs exhibited a decreased number of hair cell bundles in the lateral crista compared to controls (Figure 5C-E). Moreover, the KOs displayed a diminished acoustic startle response compared to controls (Figure 5F), suggesting a hearing impairment. Additionally, we observed reduced hair cell viability in KOs through the Yo-Pro-1 uptake experiment (Figure 5G-I). In summary, the findings of reduced otolith size, fewer hair cell bundles in the lateral crista of the inner ear, and viable hair cells in the lateral line neuromast indicating a hearing impairment in *naa80* KOs. The analysis of acoustic startle responses further substantiates these conclusions.

**Figure 5:**
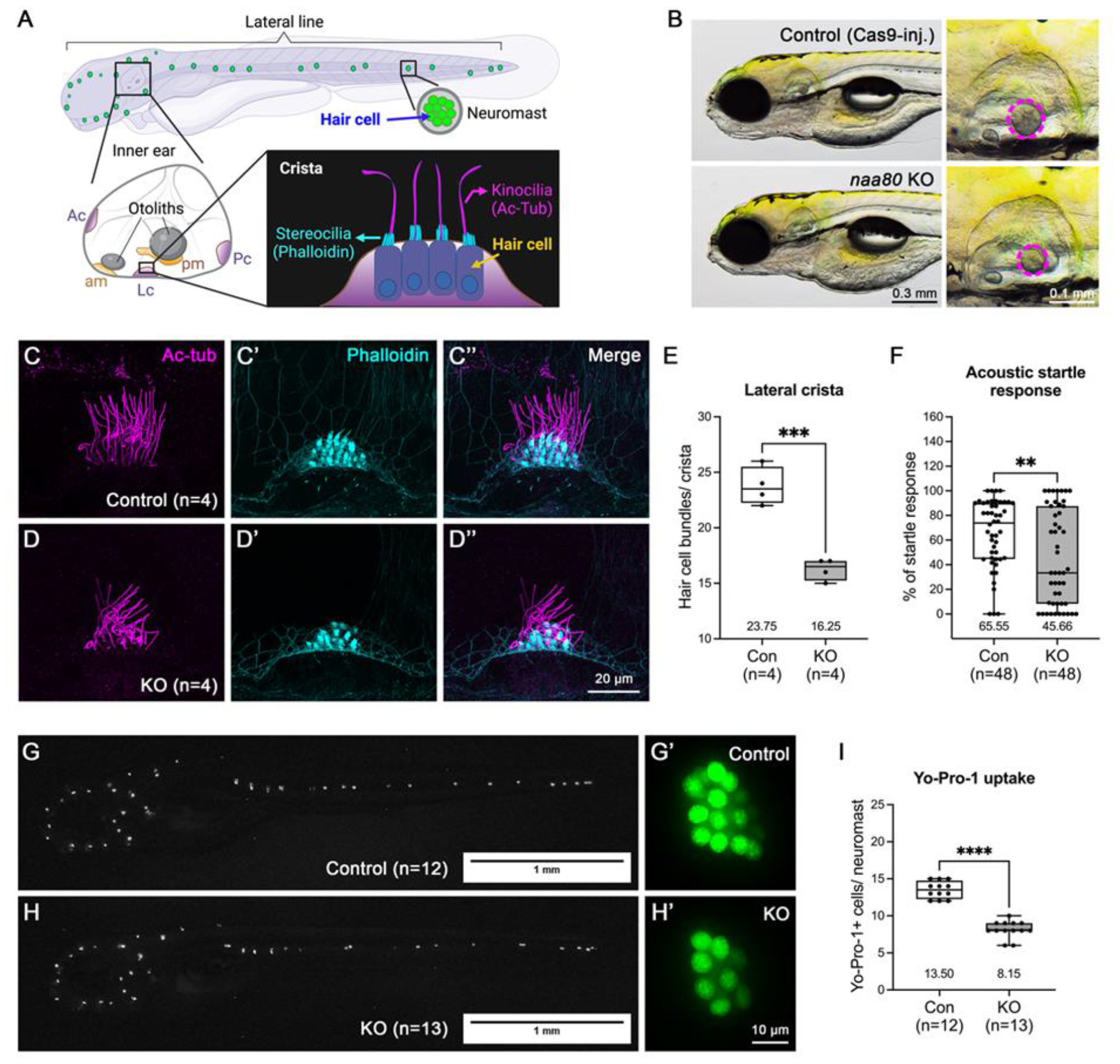
Zebrafish *naa80* F0 KO larvae showed hearing loss-related phenotypes. **A)** Schematic representation depicts sensory tissues in zebrafish larva at 5-6 dpf including inner ear and neuromasts. Five sensory patches of inner ear including anterior crista (Ac), lateral crista (Lc), posterior crista (Pc), anterior macula (am) and posterior macula (pm). Schematic representation of lateral crista at lower right corner. Kinocilia and stereocilia of hair cell can be revealed by immunohistochemistry using anti-acetylated tubulin (Ac-tub and phalloidin (F-actin), respectively. **B)** *naa80* F0 KO larvae have no obvious morphological abnormalities but showed smaller otoliths compared to Cas9-injected control larvae. **C-D)** Immunohistochemistry of both anti-acetylated tubulin (magenta, C and D) and phalloidin (cyan, C’ and D’), and merged (C’’ and D’’). **E)** Quantification of hair cell bundles in later crista. **F)** Evaluation of acoustic startle response. **G-H)** Representative image of Yo-Pro-1 uptake. **I)** Quantification of Yo-Pro-1 positive cells per neuromast. Two-tailed unpaired t test with Welch’s correction was used for E, F and I. **: *p* < 0.01, ***: *p* < 0.001 and ****: *p* < 0.0001.

## Discussion

N-terminal acetylation is probably the most common protein modification in eukaryotes since 50-90% of eukaryotic proteomes are Nt-acetylated (*23, 32, 33*). The functional impact of Nt-acetylation may vary from protein to protein and include protein folding, stability, degradation, complex formation and subcellular targeting (*34, 35*). Recent global analyses uncovered that protection from protein degradation is a major function of protein Nt-acetylation in multicellular eukaryotes (*36*–*38*). The majority of Nt-acetylation events in eukaryotes is catalyzed by a set of five conserved N-terminal acetyltransferases (NATs), NatA-NatE, acting co-translationally (*35, 39*). Furthermore, post-translational NAT enzymes were recently uncovered both in the plant and animal kingdoms (*8, 26, 40, 41*). The two currently known post-translational animal kingdom NATs are NatF/NAA60, acetylating transmembrane proteins (*25, 42*), and NatH/NAA80, acetylating actins (*8*–*10*). The NAT machinery and the Nt-acetylome have not been thoroughly studied in fish. Only basic studies on NatA and NatC have been conducted (*27, 43*). Here we investigate the mechanism and impact of Nt-acetylation of actin, the most common cellular protein in animals with numerous functions. Our results show that in zebrafish, like in humans, Nt-acetylated actin is the predominant form in wildtype fish. Estimating the Nt-acetylation stoichiometry is not possible using our methodology and would require targeted quantitation using isotopically labeled peptides, as Nt-acetyl site occupancy is close to 100% in wildtype cells (*11, 44*). Further, as we used a LysC/trypsin mix for the proteomics experiments we would not pick up N-terminally arginylated peptides if such existed (*11*), as the Nt-arginine would be removed by trypsin. Actin Nt-acetylated peptides are predominantly in the top 30^th^ percentile of peptides as ranked by intensity in *naa80*+/+ and *naa80*+/- fish and some are in the 98^th^ percentile. In the *naa80*+/+ and *naa80*+/- fish the non-Nt-acetylated actin N-termini are not found. Based on this, the Nt-acetylation stoichiometry is likely to be high. The fact that Nt-acetylated actin is undetected in all but one of the *naa80*-/- samples shows that *naa80* status is the major determinant of actin Nt-acetylation in zebrafish. The low-intensity measurement of Nt-acetylated cytoplasmic actin 2 could stem from carryover; potentially it could also be Nt-acetylated by Naa10 in *naa80*-/- fish, as zebrafish Naa10, like human NAA10, is capable of Nt-acetylating acidic N-termini without an initiator methionine (*26, 27*).

Alpha-cardiac muscle 1b actin was present with high relative intensity in the skeletal muscle samples, although conventional wisdom is that cardiac actins are mainly expressed in heart muscle. If zebrafish alpha-cardiac 1b actin (N-terminus DDEETT) is annotated as a cardiac actin simply due to similarity to human cardiac actin (N-terminal sequence is as well DDEETT (Fig. 1)) and not due to expression data, it may be the case that this isoform is indeed expressed more highly in skeletal muscle. DDDETT (alpha-skeletal 1a) is slightly higher in heart muscle, and for the same reason may be misannotated. Alternatively, but not mutually exclusively, they may both be expressed in both tissues. More focused studies should clarify this, as there has currently been little research done on actin isoforms in zebrafish.

This is the first animal knockout model of this recently characterized actin-modifying enzyme. Prior to phenotyping, several data-naïve hypotheses presented themselves. One was that actin Nt-acetylation would be so critical to the blastula-gastrula stage cell morphogenetic movements that *naa80*-/- fish would be unable to complete them. This entailed the prediction that the embryos would die during or around gastrulation, or alternatively present with large morphological abnormalities. This did not manifest, and we conclude that Naa80 is not required for normal early development.

A second but related hypothesis was that if actin Nt-acetylation is important for normal actin dynamics during the blastula stage, *naa80*-/- embryos developing from an egg laid by a *naa80*+/- mother may still have maternal *naa80* mRNA, allowing actin Nt-acetylation before the mid-blastula transition from maternal to zygotic gene expression. Maternally encoded Naa80 and Nt-acetylated actin would then be diluted into the growing embryo and eventually degraded. After the mid-blastula transition Nt-acetylated actin would not be functionally present; if actin Nt-acetylation was only required during early morphogenetic movements this could still result in a viable embryo. However, as we obtained some embryos from *naa80* 13del -/- incrosses which were viable and with no discernible morphological phenotype, we conclude that not even maternal actin Nt-acetylation is required for normal development.

A third hypothesis was that loss of actin acetylation would lead to functional changes in actin-containing structures and organs. Cultured human cells with loss of *NAA80* and actin Nt-acetylation display increased motility and increased cell size, and unacetylated actin has altered polymerization properties *in vitro* (*8*).

Muscle cells depend on actin-myosin networks to perform their contractile roles. Muscle cells lacking actin acetylation have not been described in the literature and it was unclear if such cells could perform normally. Based on our results, it appears clear that under normal conditions there is no major loss of function, as swimming and feeding behavior is not appreciably different from heterozygous or wildtype tankmates. Nt-acetylation may thus not be required for functional actin networks under normal conditions *in vivo*. Likewise, while a reasonable hypothesis was that cardiomyocytes would be impaired by loss of actin acetylation, we have no indications that heart function is diminished in *naa80* -/- zebrafish.

One drawback of our study is that we have not performed any experiments where the animals are stressed. For example, a swim-mill experiment might reveal lower endurance when the fish do not have Nt-acetylated actin. Human individuals harboring pathogenic variants of one of the actin genes display different clinical phenotypes including microcephaly, facial dysmorphism, intellectual disability, myopathy and impaired hearing (*45*–*47*). Recently, two brothers sharing the same pathogenic *NAA80* variant were identified (*16*). They both show phenotypes overlapping with those seen in individuals with impaired actin: craniofacial dysmorphisms, developmental delay, mild muscle weakness and progressive high-frequency sensorineural hearing loss. This is supportive of the main role of NAA80 in actin Nt-acetylation *in vivo* and that phenotypes driven by pathogenic *NAA80* variants may be driven by impaired actin function. Our *naa80* -/- zebrafish display reduced response to sound stimuli and actin-related hearing phenotypes in humans were particularly linked to defective γ-actin (*45*). So while many actin-related muscle functions may be normal in unstressed and unchallenged *naa80* -/- animals, Nt-acetylation could be critical in other contexts such as for non-muscle actin dynamics in the ear.

## Materials and methods

### Generation of two naa80 knockout zebrafish lines: fin clipping, gDNA isolation and genotyping

Generation of zebrafish carrying *naa80* knockout alleles was performed by the Zebrafish Genetics Core Facility, The Hospital for Sick Children, Toronto, Canada. The gRNAs for nat6/naa80 were designed using CHOPCHOP (*48*–*50*) following a previously described protocol (*51*), and synthesized using the HiScribe™ T7 High Yield RNA Synthesis Kit (NEB, E2040S), following the manufacturer’s instructions. The plasmid pT3TS-nCas9n (Addgene #46757) was used as a template to synthesize Cas9 mRNA *in vitro* using the mMESSAGE mMACHINE™ T3 Transcription Kit (Invitrogen, AM1348). To generate F0 mosaic mutant animals, 100 pg of a *naa80*-targeting gRNA (CCAATGGCAGCGCAGCATGGGGG) was injected with 150 pg Cas9 mRNA into one-cell stage AB wildtype embryos at the concentration indicated above. Adult founders carrying a 5 bp deletion/1 bp insertion (5del/1in) or 13 bp deletion (13del) were identified by Sanger sequencing using the primers ACCGTCAAAGCACATAAGAACCT and GTAGTTAGGCACAATCGGGTACA. In the CDS of the now obsolete RefSeq record XM_005167127.3 (https://www.ncbi.nlm.nih.gov/nuccore/XM_005167127.3?report=genbank), the indel positions are: 107-111 GCAGC>A for the 5del/1in line and deletion of 101-113 GGCAGCGCAGCAT for the 13del line. Each line was incrossed to obtain F1 embryos which were shipped to the zebrafish core facility in Bergen. Zebrafish were kept in the zebrafish facility at the Department of Bioscience, University of Bergen, Bergen, Norway. Fish were kept at 28.5 °C and a 14/10 hour light/dark cycle. They were fed with artemia and powdered feed twice a day. Mutant allele carriers were identified by fin clipping and sequencing. Fin clipping procedure was approved by the Norwegian Food Safety Authority (application number 20/49856, approved 15.5.2020, with changes approved 17.2.2021 and 30.6.2022).

Adult zebrafish were anesthetized by placing in 40 mg/l buffered tricaine in system water for 2-4 minutes until gill movement was reduced, placed on a petri dish, weighed, and measured.

The distal part of the tail fin was cut off with a clean scalpel, and the fish was immediately placed in system water to recover. The fish were either placed alone, with a mate of the opposite sex, or with two Spotty wild-type fish, while the genotyping was performed. gDNA was extracted from the fin essentially as described by (*52*). The fin piece was boiled for 10 minutes in 18 µl 50 mM NaOH. 2 µl 100 mM tris-HCl, pH 8.0 was added and the sample was centrifuged for 13,000 x *g* for 5 minutes. The supernatant was diluted either 1:10 or to 50 ng/µl in 100 mM tris-HCl, pH 7.4. For sequencing, 1 µl was used for a PCR reaction with primers GTAGTTAGGCACAATCGGGTACA and GTCAGCTGCACAGTCTTTCG producing a 359 bp fragment containing the mutation site. The PCR reaction ran for 35 cycles with a Tm of 58.4 °C. The PCR was checked by running on a 2% agarose/TAE gel with Gelred, and 1 µl of the PCR product was used in a Big Dye v3.1 (Applied Biosystems) sequencing reaction. Alternatively, genotyping for the 13del line was performed by agarose gel electrophoresis. 1 µl 1:10 diluted gDNA was used in a PCR reaction with primers GTAGTTAGGCACAATCGGGTACA and GGCGCTGTCTGATCTACTCC yielding a 112 bp product and a 62 °C Tm. PCR products were run on a 4% agarose gel with 10 mM lithium borate acetate (LAB) buffer and ethidium bromide, allowing the resolution of the 13 bp difference between the mutant and wildtype alleles.

### Actin Nt-acetylation status determination by mass spectrometry

Genotyped *naa80* +/+, +/- and -/- adult fish (F1) were euthanized and dissected, collecting lateral muscle, heart, and gut tissues. Fish were kept on ice after euthanasia and dissected tissue was flash-frozen in liquid nitrogen and kept at -80 °C until processing. Frozen tissues were transferred to a Petri dish on ice and chopped finely with a clean scalpel. The tissue pieces were transferred to a preweighed microcentrifuge tube and lysed in 10 µl lysis buffer per mg tissue. Lysis buffer was composed of 50 mM tris-HCl, pH 7.4, 150 mM NaCl and 1 % NP-40, supplemented with 1 tablet/50 ml of cOmplete, EDTA-free Protease Inhibitor Cocktail (Roche) and 250 U/ml benzonase. Lysis was performed for one hour on a rotating wheel at 4 °C. The lysate was cleared by centrifugation at 17,000 x *g* for 5 minutes, and the concentration was measured in the supernatant using the Pierce BCA kit according to manufacturers’ protocol. To reduce and alkylate the proteins, 50 µg of protein sample was combined with an equal volume of alkylation buffer containing 4% SDS, 10 mM TCEP, 20 mM chloroacetamide, and 200 mM tris-HCl, pH 8.0, and heated to 95 °C for 10 minutes. To digest and clean up proteins in the supernatant, we performed protein aggregation and capture (PAC) essentially as described (*53*). 250 µg Sera-Mag carboxylate-modified hydrophilic and hydrophobic magnetic beads (Cytiva) in a 50:50 ratio in acetonitrile were added to 50 µg alkylated protein, so that the final concentration of acetonitrile was 70% and the protein:bead ratio was 5:1. The samples were vortexed, left to settle for 10 minutes, vortexed and settled for 10 more minutes, and then the beads with aggregated proteins were separated from the supernatant with a magnet. The supernatant was removed with vacuum suction, and 1 ml 100% acetonitrile was added while the tubes were still on the magnet, care being taken not to disturb the beads. The acetonitrile was removed and the beads were washed on the magnet twice with 1 ml 70% ethanol. The beads were then resuspended in 250 µl digestion solution containing 1 µg trypsin and 0.5 µg LysC in 25 mM tris-HCl, pH 8.0. The beads were incubated for 16 hours at 37 °C and shaking at 1100 rpm. 1/10 volume of 10% TFA was added to each sample to acidify it and stop the digestion. The beads were separated on the magnet and the supernatant was transferred to a fresh tube, centrifuged at 13,000 x *g* for 1 minute and transferred to a new tube to remove any residual beads, and desalted using a 1 ml C18 Sep-pak (Waters) according to the manufacturer’s instructions. Peptides were dried in a speedvac, resuspended in 0.1% formic acid, and stored at -20 °C until analysis.

### LC/MS for zebrafish tissue proteomics

About 0.5 ug protein as tryptic peptides dissolved in 2% acetonitrile (ACN), 0.5% formic acid (FA), were injected into an Ultimate 3000 RSLC system (Thermo Scientific, Sunnyvale, California, USA) connected online to a Exploris 480 mass spectrometer (Thermo Fisher Scientific, Bremen, Germany) equipped with EASY-spray nano-electrospray ion source (Thermo Scientific). The sample was loaded and desalted on a pre-column (Acclaim PepMap 100, 2cm x 75µm ID nanoViper column, packed with 3µm C18 beads) at a flow rate of 5µl/min for 5 min with 0.1% trifluoroacetic acid. Peptides were separated during a biphasic ACN gradient from two nanoflow UPLC pumps (flow rate of 200 nl/min) on a 50 cm analytical column (PepMap RSLC, 50cm x 75 µm ID EASY-spray column, packed with 2µm C18 beads). Solvent A and B were 0.1% TFA (vol/vol) in water and 100% ACN respectively. The gradient composition was 5%B during trapping (5min) followed by 5-8%B over 1 min, 8–25%B for the next 124min, 25-36%B over 30 min, and 36–80%B over 5min. Elution of very hydrophobic peptides and conditioning of the column were performed during 10 minutes isocratic elution with 80%B and 15 minutes isocratic conditioning with 5%B. Instrument control was through Thermo Scientific SII for Xcalibur 1.6. The eluting peptides from the LC-column were ionized in the electrospray and analyzed by the Orbitrap Exploris 480. The mass spectrometer was operated in the DDA-mode (data-dependent-acquisition) to automatically switch between full scan MS and MS/MS acquisition. Instrument control was through Orbitrap Exploris 480 Tune 3.1 and Xcalibur 4.4. MS spectra were acquired in the scan range 350-1400 m/z with RF lens at 40%, resolution R = 120 000 at m/z 200, automatic gain control (AGC) target of 3e6 and a maximum injection time (IT) at “Auto”. The 15 most intense eluting peptides above an intensity threshold of 50 000 counts, and charge states 2 to 5 were sequentially isolated to a target value (AGC) of 1e5 and a maximum IT of 75 ms in the C-trap, and isolation width maintained at 1.2 m/z, before fragmentation in the HCD (Higher-Energy Collision Dissociation) cell. Fragmentation was performed with a normalized collision energy (NCE) of 30 %, and fragments were detected in the Orbitrap at a resolution of 30 000 at m/z 200, with first mass fixed at m/z 110. One MS/MS spectrum of a precursor mass was allowed before dynamic exclusion for 45s with “exclude isotopes” on. Lock-mass internal calibration was not used. The spray and ion-source parameters were as follows. Ion spray voltage = 1800V, no sheath and auxiliary gas flow, and capillary temperature = 275 °C.

### Database searching of proteomics data

Data was processed in Fragpipe (v. 17.1) using the LFQ-MBR workflow. The enzyme specificity was set to semi-specific trypsin (free N-terminus). The data were searched against a zebrafish proteome database containing 20,366 sequences (3,235 annotated in Swiss-Prot and 17,131 unreviewed TrEMBL sequences). Carbamidomethylation of cysteine was set as a fixed modification and N-terminal acetylation of peptide and protein N-termini as well as methionine oxidation were set as variable modifications. The combined_modified_peptide.tsv output file was used for further analysis to determine the acetylation status of N-terminal actin actin peptides.

### Generation of zebrafish founder knockout (KO) larvae for phenotyping

Two single guide RNA (sgRNA) target sequences (5’-ACTCAACATGAAAAGGTCAT-3’ with CGG PAM and 5’-TATGGCCGAATCCTTATGGA-3’ with AGG PAM) were designed using the CRISPOR tool (*54*) and chemically synthesized by Synthego Inc. The sgRNAs, along with 1 μL of 40 μM Cas9-NLS protein from UC Berkeley QB3 Macrolab (Berkeley, CA), were combined with 500 ng of each sgRNA (in 3 μL) and 2 μL of 1 M potassium chloride. This mixture was injected into one-cell stage wild-type (WT) embryos. Phenotypic analysis was conducted on the founder generation of knockouts at the 5-day post-fertilization (dpf) stage.

### Whole-mount in situ hybridization (WISH)

Whole-mount *in situ* hybridization was conducted utilizing *naa80* antisense probes labeled with digoxigenin, which were synthesized from a PCR-amplified template derived from zebrafish cDNA containing T7 promoter sequences at the 3’
s end (*55*). The specific primer sequences used were Forward 5’-GTACCCGATTGTGCCTAACT-3’ and Reverse 5’-gaattgtaatacgactcactatagggCCGAGTCTCAGATGTCCTTG-3’. RNA synthesis was performed using T7 RNA polymerase from Roche.

### RNA extraction and reverse transcription-quantitative PCR (RT-qPCR)

RNA was extracted from various developmental stages and adult tissues using TRIzol Reagent (Thermo Fischer Scientific, Waltham, MA, USA) and further purified with the RNA clean and concentrator-5 kit (Zymo), following the provided instructions. Subsequently, RNA samples were reverse transcribed into cDNA using the iScript cDNA-synthesis kit (Bio-Rad, Hercules, CA, USA), as per the manufacturer’s protocol. The generated cDNA served as a template for RT-qPCR reactions, performed with SYBR Green Supermix (Thermo Fisher Scientific) and the Light Cycler® 96 System (Roche, Pleasanton, CA) according to the manufacturer’s guidelines. Each amplification was conducted with three technical replicates and normalized to the *18S* housekeeping gene. The primer sequences used were Forward (5’-CTATCCCGTGTGTCTCCTGC-3’) and Reverse (5’-TGCAAACCACTACCGACTCC-3’).

Cycle threshold values (Ct) data were analyzed for relative gene expression using Microsoft Excel. Quantification was performed using the 2^(-ΔΔCT) method, with 18 hpf and liver samples used as calibrators for different developmental stages and adult tissues, respectively.

### Acoustic startle response assay

The Acoustic startle response test was conducted in Zebrabox behavior chambers (Viewpoint Life Sciences) at the standard room temperature, following the previously described protocol (*31*) The test assessed the percentage of responses to 12 stimuli per larva.

### Immunohistochemistry

The hair cell stereocilia in the inner ear were labeled for F-actin using fluorescently tagged (FITC) phalloidin dye, and the kinocilia were stained for anti-acetylated tubulin at 5 dpf, following the method previously outlined (*31*). Subsequently, the stained larvae were immersed in 75% glycerol, and images were captured using a Zeiss LSM-710 confocal microscope.

### Yo-Pro-1 uptake

Whole-mount staining of live zebrafish larvae was performed using Yo-Pro-1 (Invitrogen) to observe neuromast hair cells. Zebrafish larvae at 5 dpf were anesthetized with tricaine and treated with 1 mM Yo-Pro-1 diluted in 5 ml of embryo medium for 1 hour at room temperature to allow thorough probe penetration.

### Expression of zebrafish Naa80-MPB

To clone zebrafish *naa80*, a cDNA library was constructed by isolating total mRNA from a pool of 5 dpf TAB wildtype embryos essentially as described (*27*). The embryos were lysed in Trizol and RNA was extracted using chloroform/phenol followed by isopropanol precipitation. The library was generated using the Transcriptor Reverse Transcriptase kit (Roche) and poly-dT and random hexamer primers. *naa80* was amplified by PCR and subcloned into the pETM41 vector, which encodes an N-terminal His-tag and maltose binding protein (MBP) tag, ampicillin resistance, and is inducible by IPTG. pETM41-*naa80* was introduced into *Escherichia coli* BL21* cells (Invitrogen) by heatshock transformation.

Heatshocked cells were grown in rich medium for 1 hour at 250 rpm before plating on lysogeny broth-ampicillin (LB-amp) plates and overnight growth. From this we made a preculture and from this a glycerol stock, which was kept at -80 °C. The glycerol stock was sequenced to verify the plasmid sequence. To express MBP-Naa80 for purification, a preculture of 5 ml was inoculated with the glycerol stock, and the cells were grown at 250 rpm and 37 °C fro 16 hours, then diluted in a 500 ml culture and grown until they reached an optical density of 0.6 at 600 nm. Protein expression was induced by addition of 1 mM IPTG. The cells were further cultured at 20 °C and 250 rpm overnight, then harvested by centrifugation at 3095 x *g* for 30 minutes. The cell pellets were either stored at -20 °C or lysed immediately. Lysis was performed by dissolving in 10 ml ice-cold lysis buffer (20 mM imidazole, 50 mM Tris-HCl, pH 8.0, 300 mM NaCl, 1x complete EDTA-free protease inhibitor cocktail) and sonicating on ice for 7 minutes with amplitude 55 and 1 second pulses. The cell lysate was then centrifuged for 15 minutes at 17500 x *g*. The supernatant was loaded on a HisTrap 5 ml nickel-NTA column (GE Healthcare) with a peristaltic pump. The HisTrap was transferred to an ÄKTA Pure system and column was washed with 50 ml wash buffer (same as lysis buffer with no protease inhibitor) and eluted with a high imidazole buffer (300 mM imidazole, 50 mM Tris-HCl, pH 8.0, 300 mM NaCl) into a 96-deepwell plate. The relevant fractions were dialyzed into gel filtration buffer (50 mM Tris-HCl, pH 8.0, 300 mM NaCl, and 1 mM DTT. Filtered and degassed) overnight with a 12 kDa dialysis membrane, and then concentrated to a volume of 2 ml. Gel filtration chromatography was performed on the ÄKTA Pure with a Superdex 200 16/600 column and gel filtration buffer. The relevant fractions were concentrated, concentration was measured using the BCA Protein Assay Kit (Pierce) and MBP-Naa80 was diluted to 2 µM in gel filtration buffer. For long-term storage, protein was frozen in either 50% glycerol and stored at -20 °C or 10% glycerol and stored in aliquots at -80 °C. Purity was checked by SDS-PAGE.

### N-terminal acetylation assays

N-terminal acetylation activity was measured using carbon-14 labeled Ac-CoA as described (*22*). Briefly, 250 nM purified MBP-Naa80, 200 µM substrate peptide (Figure 1 and Supplemental Table S1, 50 µM ^14^C-Ac-CoA (Perkin Elmer) were mixed in acetylation buffer (50 mM Tris-HCl pH 8.5, 10% glycerol, 1mM DTT and 0.2 mM EDTA) in a 25 µl volume. Reaction proceeded for 1 hour at 37 °C. 20 µl per sample was transferred to a 1 cm^2^ piece of P81 filter paper. All pieces were washed 3 x 5 minutes in 10 mM HEPES, pH 7.4 to remove unincorporated ^14^C-Ac-CoA, and air dried. The dry filter papers were placed in scintillation vials with 5 ml Ultima Gold F scintillation cocktail (Perkin Elmer) and scintillation was measured in a TriCarb 2900TR liquid scintillation analyzer (Perkin Elmer).

### Data availability statement

All MS data are deposited to the ProteomeXchange Consortium, via the PRIDE (https://www.ebi.ac.uk/pride/) partner repository with the dataset identifier PXD046978. Username. Password: CDI6qp7C.

## Acknowledgements

Mass spectrometry-based proteomic analyses were performed by the Proteomics Unit at the University of Bergen (PROBE). This facility is a member of the National Network of Advanced Proteomics Infrastructure (NAPI), which is funded by the Research Council of Norway (INFRASTRUKTUR-program project number: 295910). This work was supported by grants from the Norwegian Health Authorities of Western Norway (F-12540 to T.A.), and the European Research Council (ERC) under the European Union Horizon 2020 Research and Innovation Program (Grant 772039 to T.A). This work was also supported by a grant from the National Institute on Deafness and Other Communication Disorders, NIH (R21DC020317 to G.K.V.), and funds from the Oklahoma Medical Research Foundation, Oklahoma City, OK, USA (G.K.V.)

## References

1. T. Svitkina, The actin cytoskeleton and actin-based motility. Cold Spring Harb. Perspect. Biol. 10, a018267 (2018).

2. T. D. Pollard, J. A. Cooper, Actin, a central player in cell shape and movement. Science 326, 1208–1212 (2009).

3. B. Bonneau, N. Popgeorgiev, J. Prudent, G. Gillet, Cytoskeleton dynamics in early zebrafish development. Bioarchitecture 1, 216–220 (2011).

4. S. Varland, J. Vandekerckhove, A. Drazic, Actin Post-translational Modifications: The Cinderella of Cytoskeletal Control. Trends Biochem. Sci. 44, 502–516 (2019).

5. J. Vandekerckhove, K. Weber, Mammalian cytoplasmic actins are the products of at least two genes and differ in primary structure in at least 25 identified positions from skeletal muscle actins. Proc. Natl. Acad. Sci. U. S. A. 75, 1106–1110 (1978).

6. P. A. Rubenstein, D. J. Martin, NH2-terminal processing of Drosophila melanogaster actin. Sequential removal of two amino acids. J. Biol. Chem. 258, 11354–11360 (1983).

7. P. A. Rubenstein, D. J. Martin, NH2-terminal processing of actin in mouse L-cells in vivo. J. Biol. Chem. 258, 3961–3966 (1983).

8. A. Drazic, H. Aksnes, M. Marie, M. Boczkowska, S. Varland, E. Timmerman, H. Foyn, N. Glomnes, G. Rebowski, F. Impens, K. Gevaert, R. Dominguez, T. Arnesen, NAA80 is actin’s N-terminal acetyltransferase and regulates cytoskeleton assembly and cell motility. Proc. Natl. Acad. Sci. U. S. A. 115, 4399–4404 (2018).

9. E. Wiame, G. Tahay, D. Tyteca, D. Vertommen, V. Stroobant, G. T. Bommer, E. Van Schaftingen, NAT6 acetylates the N-terminus of different forms of actin. FEBS J. 285, 3299–3316 (2018).

10. M. Goris, R. S. Magin, H. Foyn, L. M. Myklebust, S. Varland, R. Ree, A. Drazic, P. Bhambra, S. I. Støve, M. Baumann, B. E. Haug, R. Marmorstein, T. Arnesen, Structural determinants and cellular environment define processed actin as the sole substrate of the N-terminal acetyltransferase NAA80. Proc. Natl. Acad. Sci. U. S. A. 115, 4405–4410 (2018).

11. A. Drazic, E. Timmerman, U. Kajan, M. Marie, S. Varland, F. Impens, K. Gevaert, T. Arnesen, The Final Maturation State of β-actin Involves N-terminal Acetylation by NAA80, not N-terminal Arginylation by ATE1. J. Mol. Biol. 434, 167397 (2022).

12. H. Aksnes, M. Marie, T. Arnesen, A. Drazic, Actin polymerization and cell motility are affected by NAA80-mediated posttranslational N-terminal acetylation of actin. Commun. Integr. Biol. 11, e1526572 (2018).

13. T. B. Beigl, M. Hellesvik, J. Saraste, T. Arnesen, H. Aksnes, N-terminal acetylation of actin by NAA80 is essential for structural integrity of the Golgi apparatus. Exp. Cell Res. 390, 111961 (2020).

14. G. Rebowski, M. Boczkowska, A. Drazic, R. Ree, M. Goris, T. Arnesen, R. Dominguez, Mechanism of actin N-terminal acetylation. Sci. Adv. 6, eaay8793 (2020).

15. R. Ree, L. Kind, A. Kaziales, S. Varland, M. Dai, K. Richter, A. Drazic, T. Arnesen, PFN2 and NAA80 cooperate to efficiently acetylate the N-terminus of actin. J. Biol. Chem. 295, jbc.RA120.015468 (2020).

16. I. J. J. Muffels, E. Wiame, S. A. Fuchs, M. P. G. Massink, H. Rehmann, J. L. I. Musch, G. Van Haaften, D. Vertommen, E. van Schaftingen, P. M. van Hasselt, NAA80 biallelic missense variants result in high-frequency hearing loss, muscle weakness and developmental delay. Brain Commun. 3, fcab256 (2021).

17. P. Van Damme, M. Lasa, B. Polevoda, C. Gazquez, A. Elosegui-Artola, D. S. Kim, E. De Juan-Pardo, K. Demeyer, K. Hole, E. Larrea, E. Timmerman, J. Prieto, T. Arnesen, F. Sherman, K. Gevaert, R. Aldabe, N-terminal acetylome analyses and functional insights of the N-terminal acetyltransferase NatB. Proc. Natl. Acad. Sci. U. S. A. 109, 12449–12454 (2012).

18. P. Haahr, R. A. Galli, L. G. van den Hengel, O. B. Bleijerveld, J. Kazokaitė-Adomaitienė, J.-Y. Song, L. J. Kroese, P. Krimpenfort, M. P. Baltissen, M. Vermeulen, C. A. C. Ottenheijm, T. R. Brummelkamp, Actin maturation requires the ACTMAP/C19orf54 protease. Science 377, 1533–1537 (2022).

19. R. P. Moerschell, Y. Hosakawa, S. Tsunasawa, F. Sherman, The specificities of yeast methionine aminopeptidase and acetylation of amino-terminal methionine in vivo. J. Biol. Chem. 265, 19638–19643 (1990).

20. T. Arnesen, H. Aksnes, Actin finally matures: uncovering machinery and impact. Trends Biochem. Sci. 48, 414–416 (2023).

21. S. F. Altschul, W. Gish, W. Miller, E. W. Myers, D. J. Lipman, Basic local alignment search tool. J. Mol. Biol. 215, 403–410 (1990).

22. A. Drazic, T. Arnesen, “[14C]-Acetyl-Coenzyme A-Based In Vitro N-Terminal Acetylation Assay.” in Protein Terminal Profiling. Methods in Molecular Biology, Vol 1574, O. Schilling, Ed. (Humana Press, New York, NY, 2017).

23. T. Arnesen, P. Van Damme, B. Polevoda, K. Helsens, R. Evjenth, N. Colaert, J. E. Varhaug, J. Vandekerckhove, J. R. Lillehaug, F. Sherman, K. Gevaert, Proteomics analyses reveal the evolutionary conservation and divergence of N-terminal acetyltransferases from yeast and humans. Proc. Natl. Acad. Sci. U. S. A. 106, 8157–8162 (2009).

24. R. Evjenth, K. Hole, O. A. Karlsen, M. Ziegler, T. Amesen, J. R. Lillehaug, Human Naa50p (Nat5/San) displays both protein Nα- and Nε-acetyltransferase activity. J. Biol. Chem. 284, 31122–31129 (2009).

25. P. van Damme, K. Hole, A. Pimenta-Marques, K. Helsens, J. Vandekerckhove, R. G. Martinho, K. Gevaert, T. Arnesen, NatF contributes to an evolutionary shift in protein N-terminal acetylation and is important for normal chromosome segregation. PLoS Genet. 7, e1002169 (2011).

26. P. Van Damme, R. Evjenth, H. Foyn, K. Demeyer, P. J. De Bock, J. R. Lillehaug, J. Vandekerckhove, T. Arnesen, K. Gevaert, Proteome-derived peptide libraries allow detailed analysis of the substrate specificities of Nα-acetyltransferases and point to hNaa10p as the post-translational actin Nα-acetyltransferase. Mol. Cell. Proteomics 10, M110.004580 (2011).

27. R. Ree, L. M. Myklebust, P. Thiel, H. Foyn, K. E. Fladmark, T. Arnesen, The N-terminal acetyltransferase Naa10 is essential for zebrafish development. Biosci. Rep. 35, e00249 (2015).

28. T. Nicolson, The genetics of hearing and balance in zebrafish. Annu. Rev. Genet. 39, 9–22 (2005).

29. B. Vona, J. Doll, M. A. H. Hofrichter, T. Haaf, G. K. Varshney, Small fish, big prospects: using zebrafish to unravel the mechanisms of hereditary hearing loss. Hear. Res. 397, 107906 (2020).

30. Andreas Elepfandt, Processing of wave patterns in the lateral line system parallels to auditory processing. Acta Biol. Hung. 39, 251–65 (1988).

31. S. J. Lin, B. Vona, P. G. Barbalho, R. Kaiyrzhanov, R. Maroofian, C. Petree, M. Severino, V. Stanley, P. Varshney, P. Bahena, F. Alzahrani, A. Alhashem, A. T. Pagnamenta, G. Aubertin, J. I. Estrada-Veras, H. A. D. Hernández, N. Mazaheri, A. Oza, J. Thies, D. L. Renaud, S. Dugad, J. McEvoy, T. Sultan, L. S. Pais, B. Tabarki, D. Villalobos-Ramirez, A. Rad, J. C. Ambrose, P. Arumugam, M. Bleda, F. Boardman-Pretty, C. R. Boustred, H. Brittain, M. J. Caulfield, G. C. Chan, T. Fowler, A. Giess, A. Hamblin, S. Henderson, T. J. P. Hubbard, R. Jackson, L. J. Jones, D. Kasperaviciute, M. Kayikci, A. Kousathanas, L. Lahnstein, S. E. A. Leigh, I. U. S. Leong, F. J. Lopez, F. Maleady-Crowe, L. Moutsianas, M. Mueller, N. Murugaesu, A. C. Need, P. O’Donovan, C. A. Odhams, C. Patch, D. Perez-Gil, M. B. Pereira, J. Pullinger, T. Rahim, A. Rendon, T. Rogers, K. Savage, K. Sawant, R. H. Scott, A. Siddiq, A. Sieghart, S. C. Smith, A. Sosinsky, A. Stuckey, M. Tanguy, E. R. A. Thomas, S. R. Thompson, A. Tucci, E. Walsh, M. J. Welland, E. Williams, K. Witkowska, S. M. Wood, H. Galehdari, F. Ashrafzadeh, A. Sahebzamani, K. Saeidi, E. Torti, H. Z. Elloumi, S. Mora, T. B. Palculict, H. Yang, J. D. Wren, Ben Fowler, M. Joshi, M. Behra, S. M. Burgess, S. K. Nath, M. G. Hanna, M. Kenna, J. L. Merritt, H. Houlden, E. G. Karimiani, M. S. Zaki, T. Haaf, F. S. Alkuraya, J. G. Gleeson, G. K. Varshney, Biallelic variants in KARS1 are associated with neurodevelopmental disorders and hearing loss recapitulated by the knockout zebrafish. Genet. Med. 23, 1933–1943 (2021).

32. W. V. Bienvenut, D. Sumpton, A. Martinez, S. Lilla, C. Espagne, T. Meinnel, C. Giglione, Comparative large scale characterization of plant versus mammal proteins reveals similar and idiosyncratic N-α-acetylation features. Mol. Cell. Proteomics 11, M111.015131 (2012).

33. S. Goetze, E. Qeli, C. Mosimann, A. Staes, B. Gerrits, B. Roschitzki, S. Mohanty, E. M. Niederer, E. Laczko, E. Timmerman, V. Lange, E. Hafen, R. Aebersold, J. Vandekerckhove, K. Basler, C. H. Ahrens, K. Gevaert, E. Brunner, Identification and functional characterization of N-terminally acetylated proteins in Drosophila melanogaster. PLoS Biol. 7, e1000236 (2009).

34. H. Aksnes, R. Ree, T. Arnesen, Co-translational, Post-translational, and Non-catalytic Roles of N-Terminal Acetyltransferases. Mol. Cell 73, 1097–1114 (2019).

35. R. Ree, S. Varland, T. Arnesen, Spotlight on protein N-terminal acetylation. Exp. Mol. Med. 50, 90 (2018).

36. E. Linster, F. L. Forero Ruiz, P. Miklankova, T. Ruppert, J. Mueller, L. Armbruster, X. Gong, G. Serino, M. Mann, R. Hell, M. Wirtz, Cotranslational N-degron masking by acetylation promotes proteome stability in plants. Nat. Commun. 13, 810 (2022).

37. S. Varland, R. D. Silva, I. Kjosås, A. Faustino, A. Bogaert, M. Billmann, H. Boukhatmi, B. Kellen, M. Costanzo, A. Drazic, C. Osberg, K. Chan, X. Zhang, A. H. Y. Tong, S. Andreazza, J. J. Lee, L. Nedyalkova, M. Ušaj, A. J. Whitworth, B. J. Andrews, J. Moffat, C. L. Myers, K. Gevaert, C. Boone, R. G. Martinho, T. Arnesen, N-terminal acetylation shields proteins from degradation and promotes age-dependent motility and longevity. Nat. Commun. 14, 6774 (2023).

38. F. Mueller, A. Friese, C. Pathe, R. C. Da Silva, K. B. Rodriguez, A. Musacchio, T. Bange, Overlap of NatA and IAP substrates implicates N-terminal acetylation in protein stabilization. Sci. Adv. 7, eabc8590 (2021).

39. H. Aksnes, N. McTiernan, T. Arnesen, NATs at a glance. J. Cell Sci. 136, jcs260766 (2023).

40. W. V Bienvenut, A. Brünje, J. Boyer, J. S. Mühlenbeck, G. Bernal, I. Lassowskat, C. Dian, E. Linster, T. V Dinh, M. M. Koskela, V. Jung, J. Seidel, L. K. Schyrba, A. Ivanauskaite, J. Eirich, R. Hell, D. Schwarzer, P. Mulo, M. Wirtz, T. Meinnel, C. Giglione, I. Finkemeier, Dual lysine and N‐terminal acetyltransferases reveal the complexity underpinning protein acetylation. Mol. Syst. Biol. 16, e9464 (2020).

41. T. V. Dinh, W. V. Bienvenut, E. Linster, A. Feldman-Salit, V. A. Jung, T. Meinnel, R. Hell, C. Giglione, M. Wirtz, Molecular identification and functional characterization of the first Nα-acetyltransferase in plastids by global acetylome profiling. Proteomics 15, 2426–2435 (2015).

42. H. Aksnes, P. Van Damme, M. Goris, K. K. Starheim, M. Marie, S. I. Støve, C. Hoel, T. V. Kalvik, K. Hole, N. Glomnes, C. Furnes, S. Ljostveit, M. Ziegler, M. Niere, K. Gevaert, T. Arnesen, An organellar nα-acetyltransferase, naa60, acetylates cytosolic n termini of transmembrane proteins and maintains golgi integrity. Cell Rep. 10, 1362–1374 (2015).

43. J. M. Wenzlau, P. J. Garl, P. Simpson, K. R. Stenmark, J. West, K. B. Artinger, R. A. Nemenoff, M. C. M. Weiser-Evans, Embryonic growth - Associated protein is one subunit of a novel N-terminal acetyltransferase complex essential for embryonic vascular development. Circ. Res. 98, 846–855 (2006).

44. G. Prus, A. Hoegl, B. T. Weinert, C. Choudhary, Analysis and Interpretation of Protein Post-Translational Modification Site Stoichiometry. Trends Biochem. Sci. 44, 943–960 (2019).

45. M. Zhu, T. Yang, S. Wei, A. T. DeWan, R. J. Morell, J. L. Elfenbein, R. A. Fisher, S. M. Leal, R. J. H. Smith, K. H. Friderici, Mutations in the γ-Actin Gene (ACTG1) Are Associated with Dominant Progressive Deafness (DFNA20/26). Am. J. Hum. Genet. 73, 1082–1091 (2003).

46. J. B. Rivière, B. W. M. Van Bon, A. Hoischen, S. S. Kholmanskikh, B. J. O’Roak, C. Gilissen, S. Gijsen, C. T. Sullivan, S. L. Christian, O. A. Abdul-Rahman, J. F. Atkin, N. Chassaing, V. Drouin-Garraud, A. E. Fry, J. P. Fryns, K. W. Gripp, M. Kempers, T. Kleefstra, G. M. S. Mancini, M. J. M. Nowaczyk, C. M. A. Van Ravenswaaij-Arts, T. Roscioli, M. Marble, J. A. Rosenfeld, V. M. Siu, B. B. A. De Vries, J. Shendure, A. Verloes, J. A. Veltman, H. G. Brunner, M. E. Ross, D. T. Pilz, W. B. Dobyns, De novo mutations in the actin genes ACTB and ACTG1 cause Baraitser-Winter syndrome. Nat. Genet. 44, 440–444 (2012).

47. A. M. Kaindl, F. Rüschendorf, S. Krause, H. H. Goebel, K. Koehler, C. Becker, D. Pongratz, J. Müller-Höcker, P. Nürnberg, G. Stoltenburg-Didinger, H. Lochmüller, A. Huebner, Missense mutations of ACTA1 cause dominant congenital myopathy with cores. J. Med. Genet. 41, 842–848 (2004).

48. K. Labun, T. G. Montague, M. Krause, Y. N. Torres Cleuren, H. Tjeldnes, E. Valen, CHOPCHOP v3: Expanding the CRISPR web toolbox beyond genome editing. Nucleic Acids Res. 47, W171–W174 (2019).

49. K. Labun, T. G. Montague, J. A. Gagnon, S. B. Thyme, E. Valen, CHOPCHOP v2: a web tool for the next generation of CRISPR genome engineering. Nucleic Acids Res. 44, W272–W276 (2016).

50. T. G. Montague, J. M. Cruz, J. A. Gagnon, G. M. Church, E. Valen, CHOPCHOP: A CRISPR/Cas9 and TALEN web tool for genome editing. Nucleic Acids Res. 42, 401–407 (2014).

51. G. K. Varshney, B. Carrington, W. Pei, K. Bishop, Z. Chen, C. Fan, L. Xu, M. Jones, M. C. LaFave, J. Ledin, R. Sood, S. M. Burgess, A high-throughput functional genomics workflow based on CRISPR/Cas9-mediated targeted mutagenesis in zebrafish. Nat. Protoc. 11, 2357–2375 (2016).

52. É. Samarut, A. Lissouba, P. Drapeau, A simplified method for identifying early CRISPR-induced indels in zebrafish embryos using High Resolution Melting analysis. BMC Genomics 17, 547 (2016).

53. T. S. Batth, M. A. X. Tollenaere, P. Rüther, A. Gonzalez-Franquesa, B. S. Prabhakar, S. Bekker-Jensen, A. S. Deshmukh, J. V. Olsen, Protein aggregation capture on microparticles enables multipurpose proteomics sample preparation. Mol. Cell. Proteomics 18, 1027–1035 (2019).

54. B. González, M. Vazquez-Vilar, J. Sánchez-Vicente, D. Orzáez, “Optimization of Vectors and Targeting Strategies Including GoldenBraid and Genome Editing Tools: GoldenBraid Assembly of Multiplex CRISPR /Cas12a Guide RNAs for Gene Editing in Nicotiana benthamiana” in Recombinant Proteins in Plants. Methods in Molecular Biology, Vol 2480., S. Schillberg, H. Spiegel, Eds. (Humana, New York, NY, 2022).

55. C. Thisse, B. Thisse, High-resolution in situ hybridization to whole-mount zebrafish embryos. Nat. Protoc. 3, 59–69 (2008).

