## Supplemental Table S1 for "Naa80 is required for actin N-terminal acetylation and normal hearing in zebrafish"

**Table S1: Substrate peptides. All peptides were supplied by Innovagene with >95% purity.**

| **N-terminal sequence** | **Sequence** | **Derived from** | **Species** | **Uniprot ID** |
| --- | --- | --- | --- | --- |
| DDDIA | DDDIAALRWGRPVGRRRRPVRVYP | beta-actin | Human | P60709 |
| SESS | SESSSKSRWGRPVGRRRRPVRVYP | High mobility group protein A1 | Human | P17096 |
| MELL | MELLSPPRWGRPVGRRRRPVRVYP | MyoD | Human | P15172 |
| MLGP | MLGPEGG RWGRPVGRRRRPVRVYP | hnRNP F | Human | P52597 |
| MAPL | MAPLDLD RWGRPVGRRRRPVRVYP | Protein phosphatase 6 | Human | O00743 |
| DDEI | DDEIAALRWGRPVGRRRRPVRVYP | Actin, cytoplasmic 2 | Zebrafish | Q7ZVF9 |
| DEEI | DEEIAALRWGRPVGRRRRPVRVYP | Actin, cytoplasmic 1 | Zebrafish | Q7ZVI7 |
| DDDE | DDDETTARWGRPVGRRRRPVRVYP | Alpha-skeletal muscle 1a | Zebrafish | F1QUN8 |
|  |  | Alpha-skeletal muscle 1b | Zebrafish | Q4KMI7 |
| DDEES | DDEESTARWGRPVGRRRRPVRVYP | Alpha-smooth muscle 2 | Zebrafish | Q6DHS1 |
| DDEET | DDEETTARWGRPVGRRRRPVRVYP | Alpha-cardiac muscle 1b | Zebrafish | Q9I8V1 |
| DEEE | DEEETTARWGRPVGRRRRPVRVYP | Alpha-cardiac muscle 1a | Zebrafish | F1RCB6 |
